# Accelerated evolution of oligodendrocytes in human brain

**DOI:** 10.1101/601062

**Authors:** Stefano Berto, Isabel Mendizabal, Noriyoshi Usui, Kazuya Toriumi, Paramita Chatterjee, Connor Douglas, Carol Tamminga, Todd M. Preuss, Soojin V. Yi, Genevieve Konopka

**Author notes:** Correspondence (G.K.), (S.V.Y.). Co-first author (Stefano Berto, Isabel Mendizabal, Noriyoshi Usui, Kazuya Toriumi).

## Abstract

Recent discussions of human brain evolution have largely focused on increased neuron numbers and changes in their connectivity and expression. However, it is increasingly appreciated that oligodendrocytes play important roles in cognitive function and disease. Whether both cell-types follow similar or distinctive evolutionary trajectories is not known. We examined the transcriptomes of neurons and oligodendrocytes in the frontal cortex of humans, chimpanzees, and rhesus macaques. We identified human-specific trajectories of gene expression in neurons and oligodendrocytes and show that both cell-types exhibit human-specific upregulation. Moreover, oligodendrocytes have undergone accelerated gene expression evolution in the human lineage compared to neurons. The signature of acceleration is enriched for cell type-specific expression alterations in schizophrenia. These results underscore the importance of oligodendrocytes in human brain evolution.

## INTRODUCTION

Increased brain size, accompanied by increased neuron numbers, has been a central theme in human brain evolutionary studies (Gabi et al., 2016; Preuss, 2017). However, such changes alone are unlikely to entirely account for the evolved cognitive capabilities of humans (Sousa et al., 2017a). Changes in gene expression have been hypothesized as a key facet of human brain evolution (Khaitovich et al., 2006; King and Wilson, 1975), and previous bulk transcriptome studies have shown that gene expression changes in neurons have been extensive (Konopka et al., 2012; Liu et al., 2012; Sousa et al., 2017b). However, non-neuronal cell-types, particularly oligodendrocytes, show altered functional and disease-related patterns in humans compared to other primates (Donahue et al., 2018; Miller et al., 2012; Rilling and van den Heuvel, 2018). For example, compared to non-human primates, human brains have greater than expected connectivity requiring myelination (Rilling and van den Heuvel, 2018), myelination in human brains has a protracted developmental timing, and myelination and oligodendrocyte function has been implicated in neuropsychiatric diseases such as schizophrenia (SZ) (Mighdoll et al., 2015). Moreover, ∼75% of non-neurons in the human cortex consist of oligodendrocytes (Herculano-Houzel, 2014; Pelvig et al., 2008). Along with the growing appreciation of oligodendrocyte involvement in cognition (Fields et al., 2014; Voineskos et al., 2013), this suggests that oligodendrocytes may have been important targets of change in human brain evolution.

## RESULTS

### Cell-type evolutionary trajectories highlight oligodendrocyte acceleration on the human lineage

To address the contribution of cell-types to human brain evolution, we compared the cell-type specific transcriptome profiling of sorted nuclei from humans to chimpanzees, our closest extant relative, using rhesus macaque as an outgroup. We analyzed genome-wide expression levels in adult human Brodmann area 46 (BA46, NeuN: n=27, OLIG2: n=22), and the homologous regions of chimpanzee (NeuN: n=11, OLIG2: n=10), and rhesus macaque (NeuN: n=15, OLIG2: n=10) (Table S1). Prefrontal area BA46 was selected due to its association with human-specific cognitive abilities and evolution as well as neuropsychiatric disorders (Donahue et al., 2018; Fu et al., 2011; Teffer and Semendeferi, 2012). Cell-type specific whole transcriptome data was obtained using fluorescence-activated nuclei sorting (FANS) (Jiang et al., 2008) with antibodies to either NeuN or OLIG2 to isolate neurons (NeuN) or oligodendrocytes and their precursors (OLIG2), respectively (Figure S1A-E). Covariates such as age and sex, and technical confounders such as RNA integrity number (RIN) explained only a small portion of the variance in both cell-types (Figure S1F). Comparisons of our data with single-cell transcriptome data from human brain (Boldog et al., 2018) demonstrate that NeuN gene expression was representative of both inhibitory and excitatory neuronal expression signatures while OLIG2 gene expression was primarily representative of oligodendrocyte expression signatures, supporting our FANS approach (Figure S1G).

Using only high-confidence orthologous genes, we detected 8759 protein-coding genes expressed in at least one species in NeuN and 7362 protein-coding genes in OLIG2. Principal component analysis revealed that gene expression in each cell type separated by species (Figure 1A-B). Using a parsimony method, we detected species-specific differentially expressed genes (DEG) (Figure S1H; Table S2; Methods). The lineage connecting the rhesus macaque to the ancestor of humans and chimpanzees had the largest number of DEGs, which is consistent with the idea that gene expression changes accumulate with divergence times (Figure 1C) (Konopka et al., 2012; Liu et al., 2012; Sousa et al., 2017b). Furthermore, greater number of genes exhibited upregulation compared to downregulation in the human lineage, for both NeuN and OLIG2 in comparison with the non-human primates (*X*^2^ test, p= 1×10^-06^ and p = 4×10^-05^ respectively) (Figure 1C). Interestingly, human-specific expression was more pronounced in OLIG2 compared to NeuN. Specifically, OLIG2 exhibited greater effect sizes in pairwise comparisons (human versus chimpanzee, mean(log _2_(FC)): 0.26 NeuN, 0.59 OLIG2, *P* < 2 × 10^-16^, K-S test; human versus rhesus macaque, mean(log _2_(FC)): 0.25 NeuN, 0.58 OLIG2, *P* < 2 × 10^-16^, K-S test) as well as greater number of differentially expressed genes per million years (Figure 1D-E) compared to NeuN. Importantly, to ensure that these observations were not due to different numbers of samples in each species, we used a cross-validation method by using the same numbers of samples between species, which effectively is a downsampling of the human and macaque samples (Figure S2; Methods). Doing so reduced the number of DEGs per million year due to smaller sample size and heterogeneity within and between species; however, downsampled DEGs show the same pattern of acceleration in OLIG2 compared with NeuN, further supporting the result that oligodendrocytes have undergone an evolutionary acceleration on the human lineage. We next examined previous gene expression studies of frontal cortex evolution (Berto and Nowick, 2018; Konopka et al., 2012; Somel et al., 2010; Sousa et al., 2017b) to assess how bulk tissue expression profiles may have been confounded by cell-type specific trajectories. We found that human-specific upregulated genes in the bulk tissue studies were enriched with human-specific NeuN upregulated DEGs. In comparison, human-specific DEGs in OLIG2 were not enriched in the previous studies (Figure 1F), indicating that studies using bulk tissues may have been underpowered to detect oligodendrocyte-specific evolutionary trajectories. Carrying out deconvolution analysis, we found that these bulk RNA-seq datasets were primarily comprised of neuronally derived gene-expression signatures (Figure S3A-D). Thus, using a cell-type specific approach, we detected a previously undiscovered signal of rapid acceleration of oligodendrocyte-gene expression compared with neurons in the human lineage.

**Figure 1.**
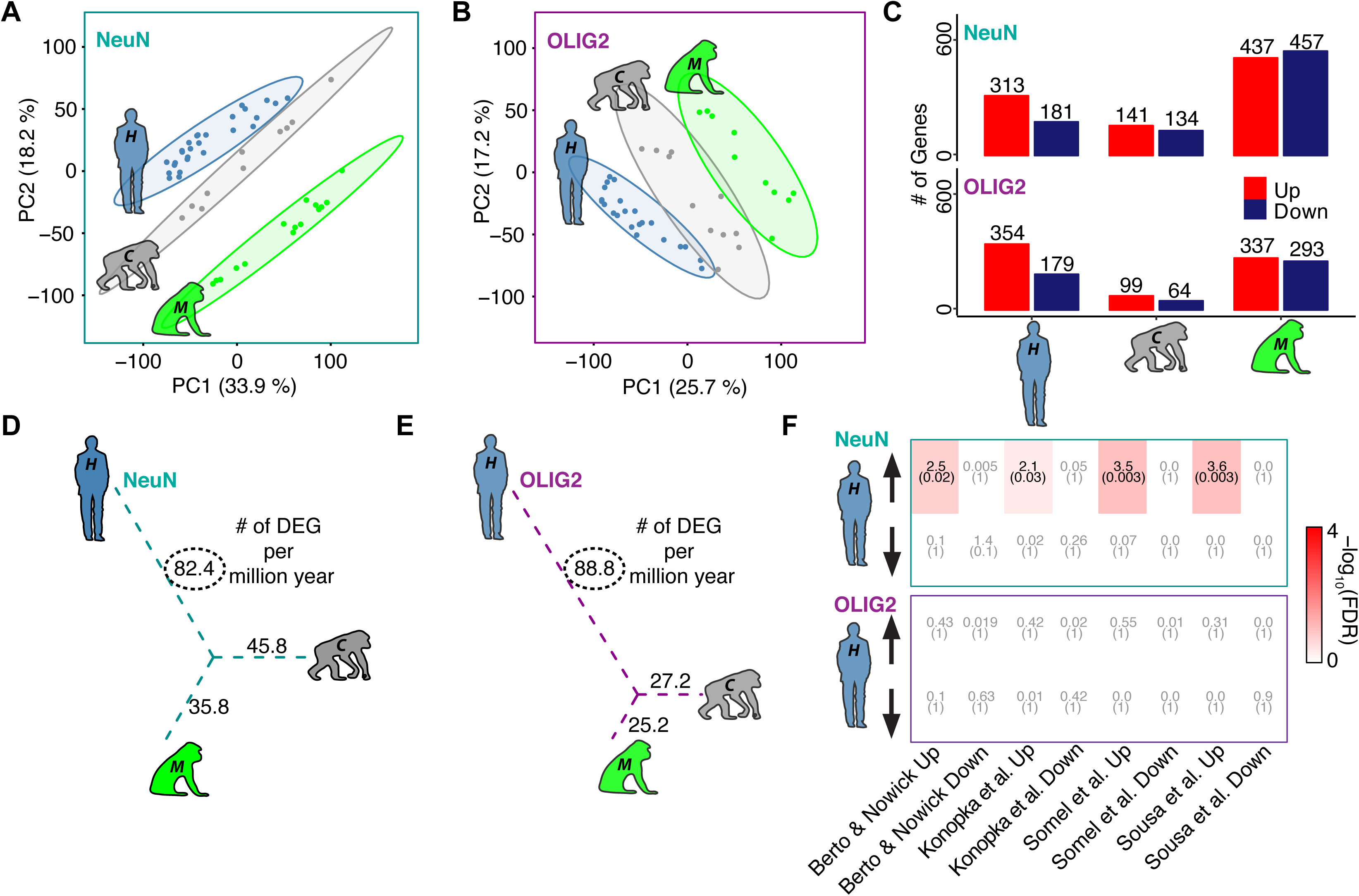
Cell type-specific differential gene expression analysis of three primates. A-B) Principal component analysis of NeuN (A) and OLIG2 (B) nuclei. Blue = human, grey = chimpanzee, green = rhesus macaque. C) Barplots representing species-specific differentially expressed genes divided by up- (red) and downregulated (darkblue) for both NeuN and OLIG2. D-E) The number of DEGs per million year (myr; human = 6 myr, chimpanzee = 6 myr, rhesus macaque = 25 myr) for the unrooted tree of the study species of NeuN (D) and OLIG2 (E). (F) Heatmap showing FDR (parenthesis) and Odds Ratio of gene set enrichment for both for both NeuN (top) and OLIG2 (bottom). Enrichment is based on a Fisher’s exact test. X-axis shows the representative data included for this analysis (Berto and Nowick, 2018; Konopka et al., 2012; Somel et al., 2010; Sousa et al., 2017b).

### Gene co-expression network highlights human-specific modules

To place the human specific changes within a systems-level context and identify the relevant biological processes associated with these changes, we next applied a permuted weighted gene co-expression analysis (Langfelder and Horvath, 2008) to detect human-specific cell-type co-expression modules (Figure 2A; Table S3; Methods). Using the expression data that were adjusted for potential variation explained by covariates and surrogate variables, we defined two modules in NeuN and two modules in OLIG2 that exhibited human-specific expression and showed a strong enrichment for human-specific DEGs (Figure 2B and C; Figure S4A-B). These modules showed higher association with species than with other covariates (Figure S4C-D). Human-specific DEGs in both NeuN and OLIG2 exhibited significantly greater connectivity compared with other genes across all modules, indicating their pivotal roles in human frontal cortex transcriptional networks (Figure 2D). NeuN human upregulated module NM21 was enriched for genes involved in synaptic function and vesicular transport (Figure 2E; Table S3). Interestingly, the OLIG2 human upregulated module OM15 was enriched for pathways implicated in RNA splicing, RNA metabolism, and chromatin remodeling (Figure 2F; table S3). Whereas downregulated module NM19 was not enriched for any specific function (Figure 2G), the OLIG2 downregulated module OM2 functions were related to transcriptional regulation, histone methylation and modification (Figure 2H; Table S3). Of note, both OLIG2 modules that are associated with human-specific expression are significantly enriched for transcription factors and RNA binding proteins (Figure S4E-F). These results suggest that alternative splicing and transcriptional regulation are biological functions linked with oligodendrocyte evolution in the human frontal cortex.

**Figure 2.**
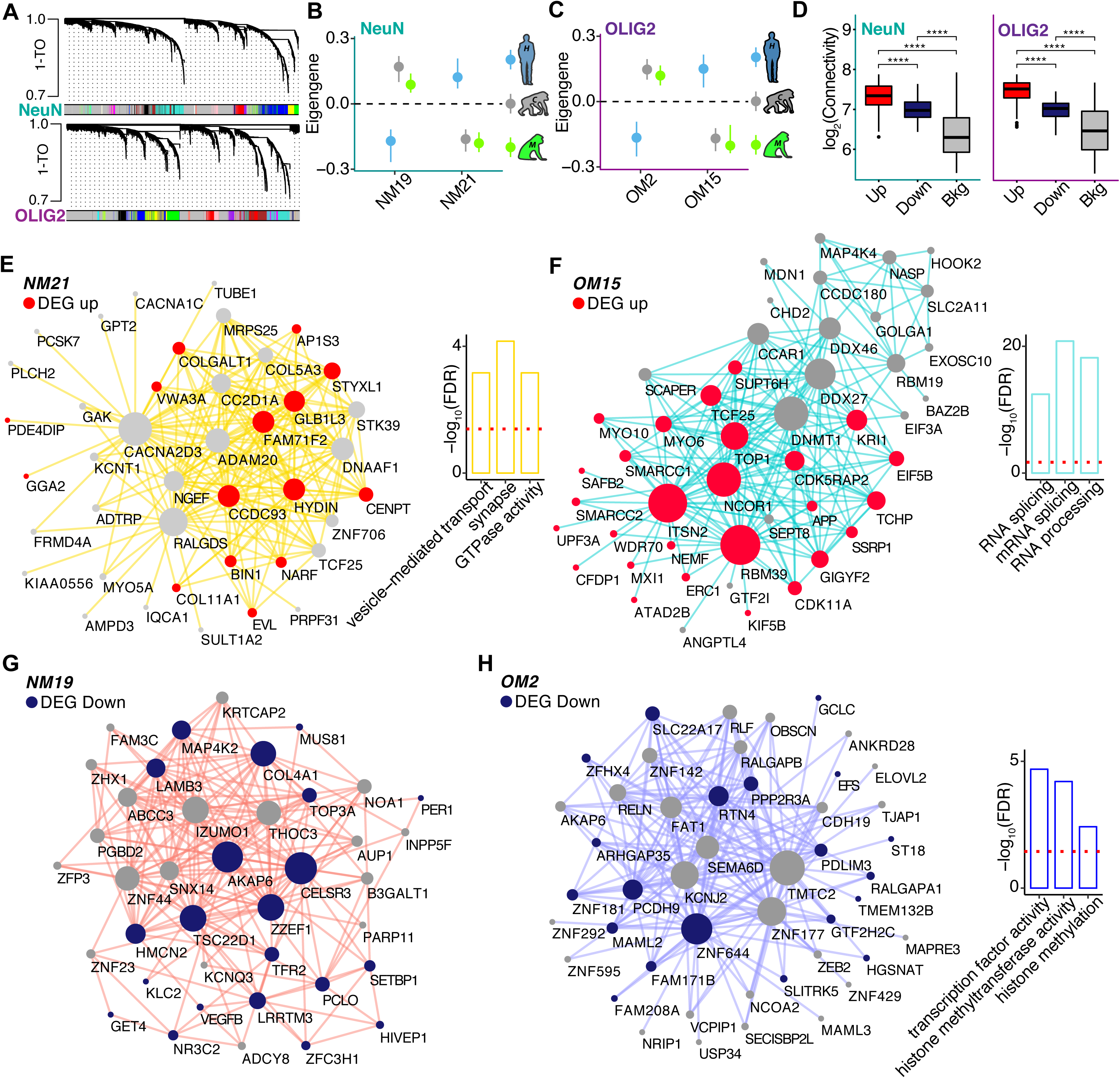
Co-expression analyses identify human-specific modules. A) Representative network dendrograms for NeuN (top) and OLIG2 (bottom). B-C) Dotplots with standard errors demonstrate the association of the modules detected by parsimony with species for NeuN (B) and OLIG2 (C). Standard errors are calculated based on the eigengene across samples. Dots represent the mean eigengene for that module. (D) Boxplots show the difference in connectivity between human-specific genes in NeuN (left) and OLIG2 (right) across the entire co-expression network compared with the background genes (**** = P < 0.001; Wilcoxon’s rank sum test). E-H) Visualization of the top 200 connections ranked by weighted topological overlap values for NM21 (E), OM15 (F), NM19 (G), and OM2 (H). Node size corresponds to the number of edges (*degree*). Human-specific upregulated genes are highlighted in red. Human-specific downregulated genes are highlighted in blue. Side barplots show the top three functions of the module based on –log _10_(FDR). Red dashed line corresponds to the FDR threshold of 0.05.

### Oligodendrocyte human-specific modules are enriched for variants associated with neuropsychiatric disorders

It is hypothesized that genes important for the evolution of human-specific cognitive abilities are linked with human-specific cognitive disorders (Doan et al., 2018; Hardingham et al., 2018). To investigate this hypothesis, we assessed the enrichment of human-specific co-expression modules with GWAS signals (Methods). While we did not find any enrichment for neuropsychiatric disorders GWAS signals in the human-specific neuronal modules NM19 and NM21 (Figure 3A), the OLIG2 human downregulated module OM2 showed a strong enrichment for attention deficit hyperactivity disorder (ADHD), bipolar disorder (BD), and SZ loci as well as loci associated with cognitive traits, education attainment and intelligence (Figure 3B; Table S4). The upregulated module OM15 showed enrichment for major depressive disorder (MDD) (Figure 3B; Table S4). In comparison, neither NeuN nor OLIG2 human-specific modules showed significant enrichment for GWAS signals associated with non-brain related traits/disease. While very little is known about the role of oligodendrocytes in ADHD (Dark et al., 2018), BD, SZ, and MDD have been associated with alterations in white matter and differential regulation of oligodendrocyte-related genes (Barley et al., 2009; Haroutunian et al., 2014; Srivastava et al., 2016; Tonnesen et al., 2018). These results suggest that human-specific co-expressed genes that are under evolutionary expression trajectories in oligodendrocytes are at risk to be associated with cognitive disease-related variants.

**Figure 3.**
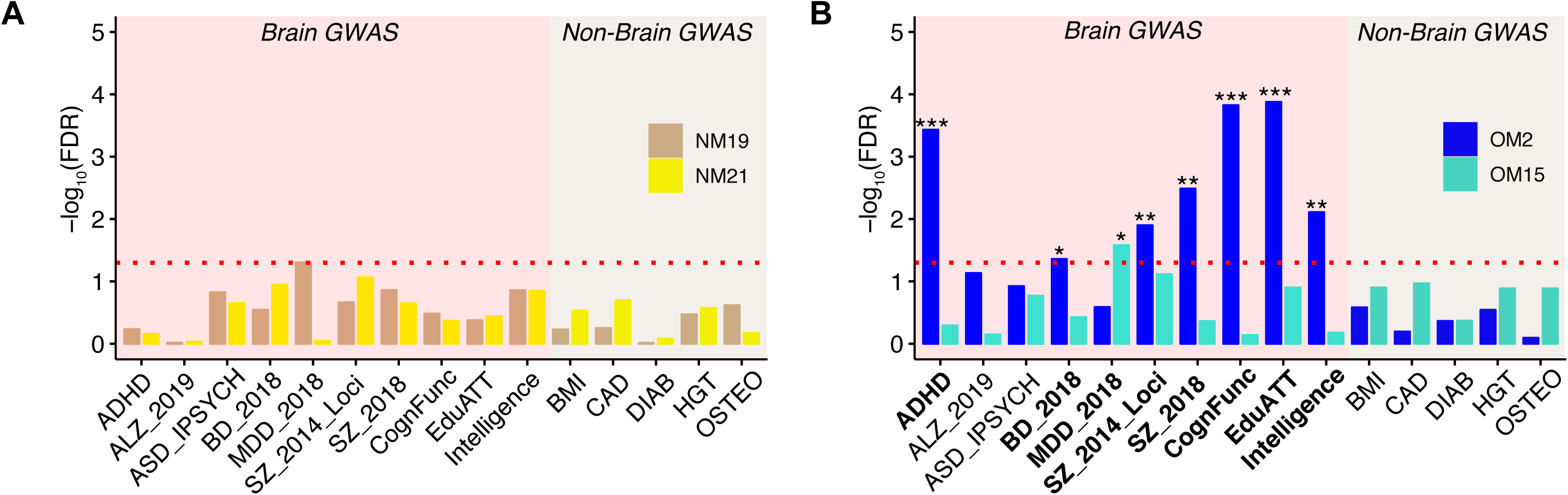
Human-specific genes are enriched for cognitive disease risk variants. A-B) Barplots highlighting the enrichment for genetic variants (–log _10_(FDR)). Bars correspond to (A) NeuN modules (NM19: tan, NM21: yellow) and (B) OLIG2 modules (OM2: blue, OM15: turquoise) species-specific modules (*** = FDR < 0.001, ** = FDR < 0.01, * = FDR < 0.05; MAGMA statistics). Red dashed line corresponds to the FDR threshold of 0.05. X-axis shows the acronyms for the GWAS data utilized for this analysis. *ADHD: attention deficit hyperactivity disorder, ALZ: Alzheimer’s disease, ASD: autism spectrum disorder from IPSYCH (Integrative Psychiatric Research), BP: bipolar disorder, MDD: major depressive disorder, SZ: schizophrenia, CognFunc: Cognitive functions, EduATT: educational attainment, BMI: body mass index, CAD: coronary artery disease, DIAB: diabetes, HGT: Height, OSTEO: osteoporosis.*

### Human-specific modules are enriched for neuropsychiatric differentially expressed genes

To further examine the potential relationship between dysregulation in neuropsychiatric disorders and human-specific changes using a large-scale gene expression dataset, we used recently published meta-analyses of cognitive disease brain gene expression from the PsychENCODE Consortium (Gandal et al., 2018) (Methods). We found that the NeuN human-specific upregulated module NM21 is overrepresented for genes in a neuronal module dysregulated in SZ and autism spectrum disorder (ASD) (geneM8; OR = 9.7, FDR = 1 × 10^-08^; Figure 4A). In contrast, the OLIG2 human-specific downregulated module OM2 is enriched for genes in an oligodendrocyte module containing genes dysregulated in ASD, BD, and SZ (geneM2; OR = 9.5, FDR = 8 × 10^-09^; Figure 4B). Interestingly, the OLIG2 upregulated module OM15 is enriched for genes in a module dysregulated in SZ and linked with splicing (geneM19; OR=10.1, FDR=1 × 10^-09^; Figure 4B), reflecting the functional enrichment we described (Figure 2F). We next assessed whether human-specific cell-type expression patterns are at risk in neuropsychiatric disorders using cell type-specific disease-relevant gene expression data (Methods). We examined cell type-specific whole transcriptome data from BA46 from 23 patients with SZ, generated following identical experimental procedures (unpublished data). Using genes differential expressed between SZ and healthy donors at the cell-type level (referred to as ‘szDEGs’; Table S5), we asked whether dysregulated genes in SZ were enriched for human-specific evolutionary changes of gene co-expression at the cell-type level. Whereas NeuN modules were not found enriched for cell-type SZ genes (Figure 4C), we found that OLIG2 szDEGs were enriched for human OLIG2 modules (Figure 4D). Specifically, the downregulated module OM2 is enriched for SZ OLIG2 upregulated genes (OR = 2.9, FDR = 5 × 10^-04^) while the upregulated module OM15 is moderately enriched for SZ OLIG2 downregulated genes (OR = 1.6, FDR = 0.05). Taken together, these observations highlight the link between oligodendrocyte evolution and neuropsychiatric disease etiologies.

**Figure 4.**
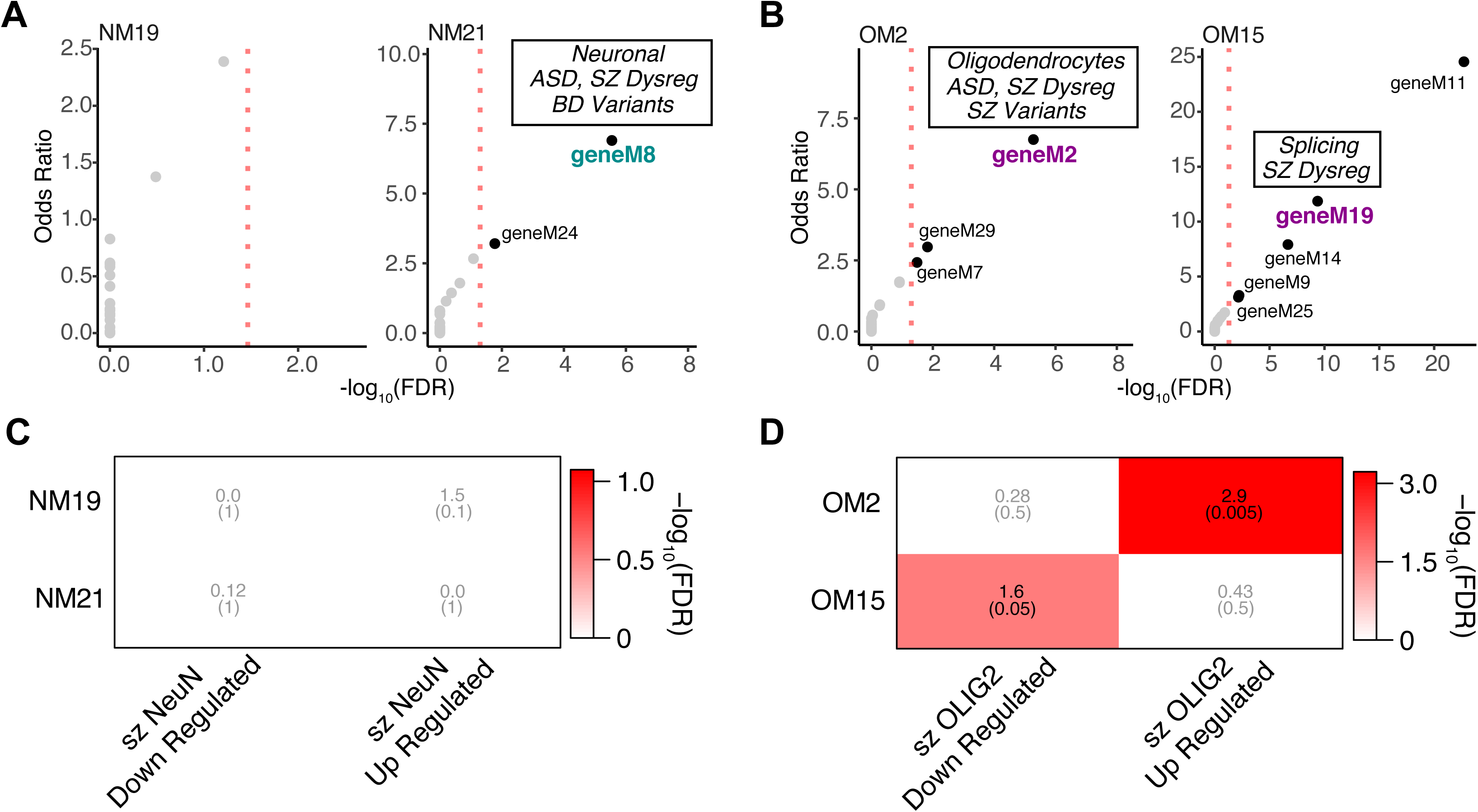
Cell type-specific expression in schizophrenia is related to human-specific genes. A) Bubble chart illustrates -log _10_(FDR) (X-axis) and Odds Ratio (Y-axis) of gene set enrichment for gene modules implicated in neuropsychiatric disorders (Gandal et al., 2018) and NeuN human-specific modules. Marked the modules with functional conservation. *Dysreg: dysregulated.* B) Bubble chart illustrates -log _10_(FDR) (X-axis) and Odds Ratio (Y-axis) of gene set enrichment for gene modules implicated in neuropsychiatric disorders (Gandal et al., 2018) and OLIG2 human-specific modules. The modules with functional conservation with the PsychENCODE dataset are indicated. C) Heatmap illustrates FDR (parenthesis) and Odds Ratio of gene set enrichment (Fisher’s exact test). X-axis shows NeuN human-specific up-/downregulated genes. Y-axis shows SZ differentially expressed genes up-/downregulated in NeuN. D) Heatmap illustrates FDR (parenthesis) and Odds Ratio of gene set enrichment. X-axis shows OLIG2 human-specific up-/downregulated. Y-axis shows SZ differentially expressed genes up-/downregulated in OLIG2.

## DISCUSSION

These data provide novel insights into cell-type species-specific expression patterns during primate brain evolution. Much of the recent focus on human brain evolution has highlighted changes in neuronal number and function in the human brain; however, the molecular characterization of the mechanisms driving such changes in neurons and as well as other cell-types is critical for understanding human brain evolution. Here, we show that NeuN human*-*specific DEGs encode genes important for synaptic function in line with previous data from bulk RNA-seq (Konopka et al., 2012; Liu et al., 2012; Sousa et al., 2017b). Surprisingly though, we find that gene expression in oligodendrocytes has undergone a more dramatic acceleration on the human lineage compared with neurons. We also show that previous comparative primate gene expression studies were likely underpowered to detect these non-neuronal expression changes.

The human-specific oligodendrocyte genes are enriched for functional categories such as RNA metabolism and RNA processing. While such molecular functions are underexplored with respect to oligodendrocytes, there is increasing evidence that these functions are altered in cognitive diseases (Glatt et al., 2011; Quesnel-Vallieres et al., 2018; Reble et al., 2018). Moreover, the brain GWAS and PsychENCODE enrichments for cell-type expression modules suggest that human-specific cell type-specific evolutionary trajectories of gene expression are implicated in disease pathophysiology in multiple cognitive disorders. Using the only available disease cell-type expression dataset, we also observe SZ cell type-specific downregulation among human-specific oligodendrocyte genes. Together, these data suggest a role for human-specific oligodendrocyte genes in disorders including SZ, MDD, BD, and ADHD. Previous work has specifically singled out oligodendrocyte dysfunction in both SZ and MDD (Miyata et al., 2015). In addition, human brains have undergone a volumetric expansion of white matter (Donahue et al., 2018; Rilling and van den Heuvel, 2018), while these white matter volumes are significantly reduced in SZ (Davis et al., 2003; Mighdoll et al., 2015). While we have focused on the novelty of the human-specific oligodendrocyte genes, there are clearly evolutionarily relevant changes in neurons that are likely important for cognitive disorders such as SZ and ASD too. Since neuronal activity can direct oligodendrocyte development and myelination (Gibson et al., 2014), the functional outcome of the interplay of gene expression changes in these two cell-types may be important for multiple cognitive disorders. Future studies that connect changes in the functional properties of oligodendrocytes, for example at the level of RNA binding and/or processing, to either disease pathophysiology, white matter volume alterations, or response to neuronal activity will confirm the importance of the identified genes in human brain evolution. Our study highlights the importance of non-neuronal cell-types in brain evolution and cognitive disorders.

## Figure Legends

**Figure S1.**
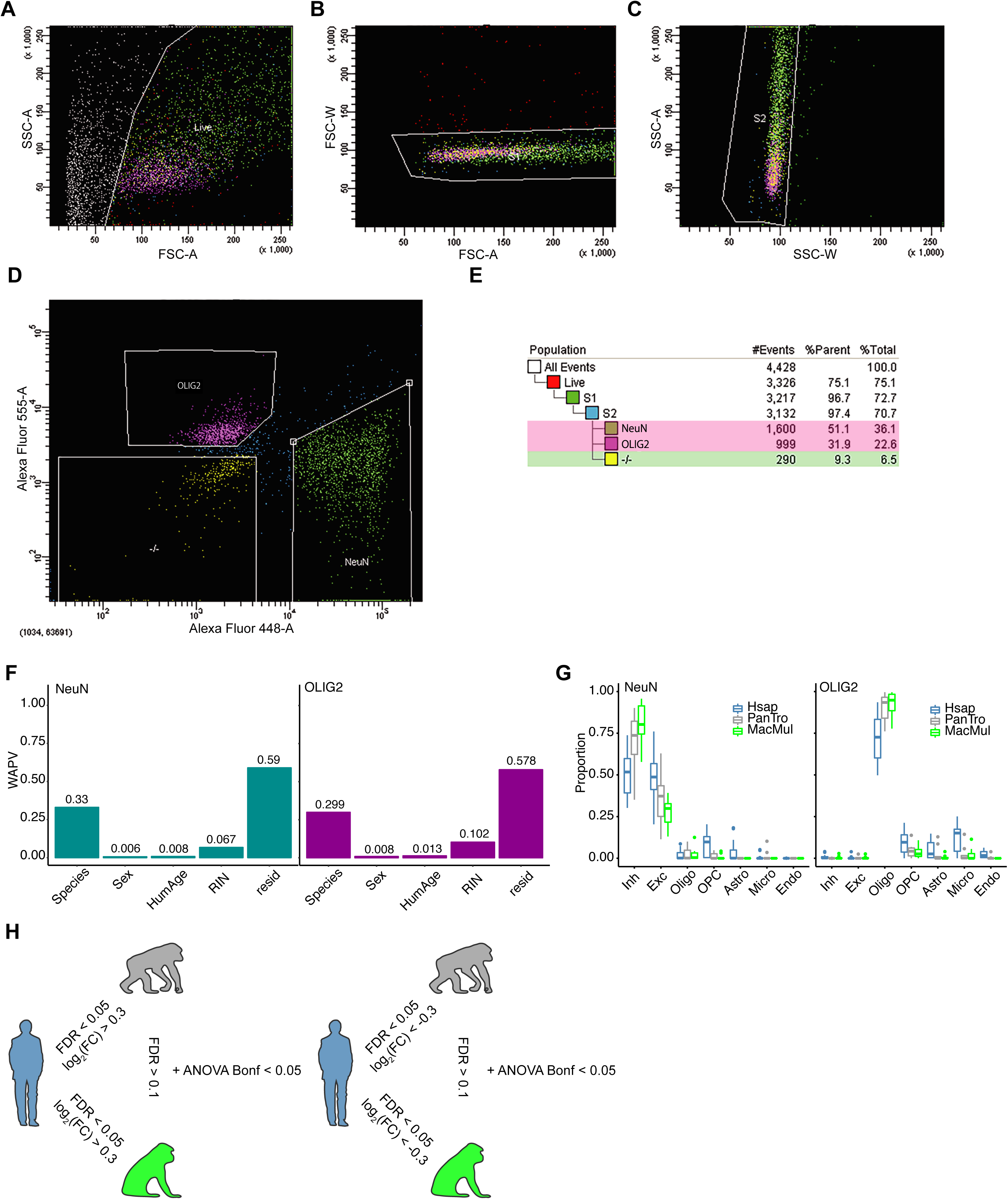
Generation and analyses of human-specific cell-type gene expression profiles. A-E) An example isolation of nuclei expressing either NeuN conjugated to Alexa 488 or OLIG2 conjugated to Alexa 555. Nuclei were first sorted for size and complexity for removing dead cells (A), followed by gating to exclude doublets that indicate aggregates of nuclei B-C), and then further sorted to isolate nuclei based on fluorescence (D). “-/-” nuclei are those that are neither NeuN+ nor OLIG2+. (E) An example of percentage nuclei at each selection step during FANS of a chimpanzee sample. Note that while in this example more nuclei were NeuN+, in other samples, the proportions might be reversed. NeuN-positive (NeuN+) nuclei represent neurons within the cerebral cortex as few NeuN-negative (NeuN-) cells in the mammalian cortex are neurons (e.g. Cajal-Retzius neurons). OLIG2-positive (OLIG2+) nuclei represent oligodendrocytes and their precursors. F) Variance explained by covariates weighted across the first 5 principal components (WAPV = Weighted average proportion variance) for NeuN or OLIG2. Gene expression patterns showed small variance explained by biological (Sex and Humanized Age) and technical (RIN) covariates. G) Deconvolution based on Allen Brain Institute single nuclei data from the human middle temporal gyrus (MTG) (Boldog et al., 2018). NeuN is explained by a high proportion of inhibitory and excitatory neurons. OLIG2 is explained by oligodendrocytes. Y-axis represents weighted proportion. X-axis represents the cell-type identified in the Allen Brain Institute single nuclei data from MTG (Hsap = *Homo sapiens*, PanTro = *Pan troglodytes*, MacMul = *Macaca mulatta*, Inh = inhibitory neurons, Exc = excitatory neurons, Oligo = oligodendrocytes, OPC = oligodendrocyte precursor cells, Astro = astrocytes, Micro = microglia, Endo = endothelial cells). H) A parsimony approach based on linear model statistics from pairwise comparisons. Left panel: Upregulated genes were considered if FDR < 0.05 in species 1 vs species 2 and species 1 vs species 3 with log _2_(FC) > 0.3 in both comparisons while not significantly differentially expressed (defined as FDR > 0.1) in species 2 vs species 3. Similarly, right panel: Downregulated genes were considered if FDR < 0.05 in species 1 vs species 2 and species 1 vs species 3 with log _2_(FC) < -0.3 in both comparisons with FDR > 0.1 in species 2 vs species 3. Additionally, we added a Bonferroni corrected ANOVA < 0.05 cutoff in the model across the three species analyzed. Blue = human, grey = chimpanzee, green = rhesus macaque.

**Figure S2.**
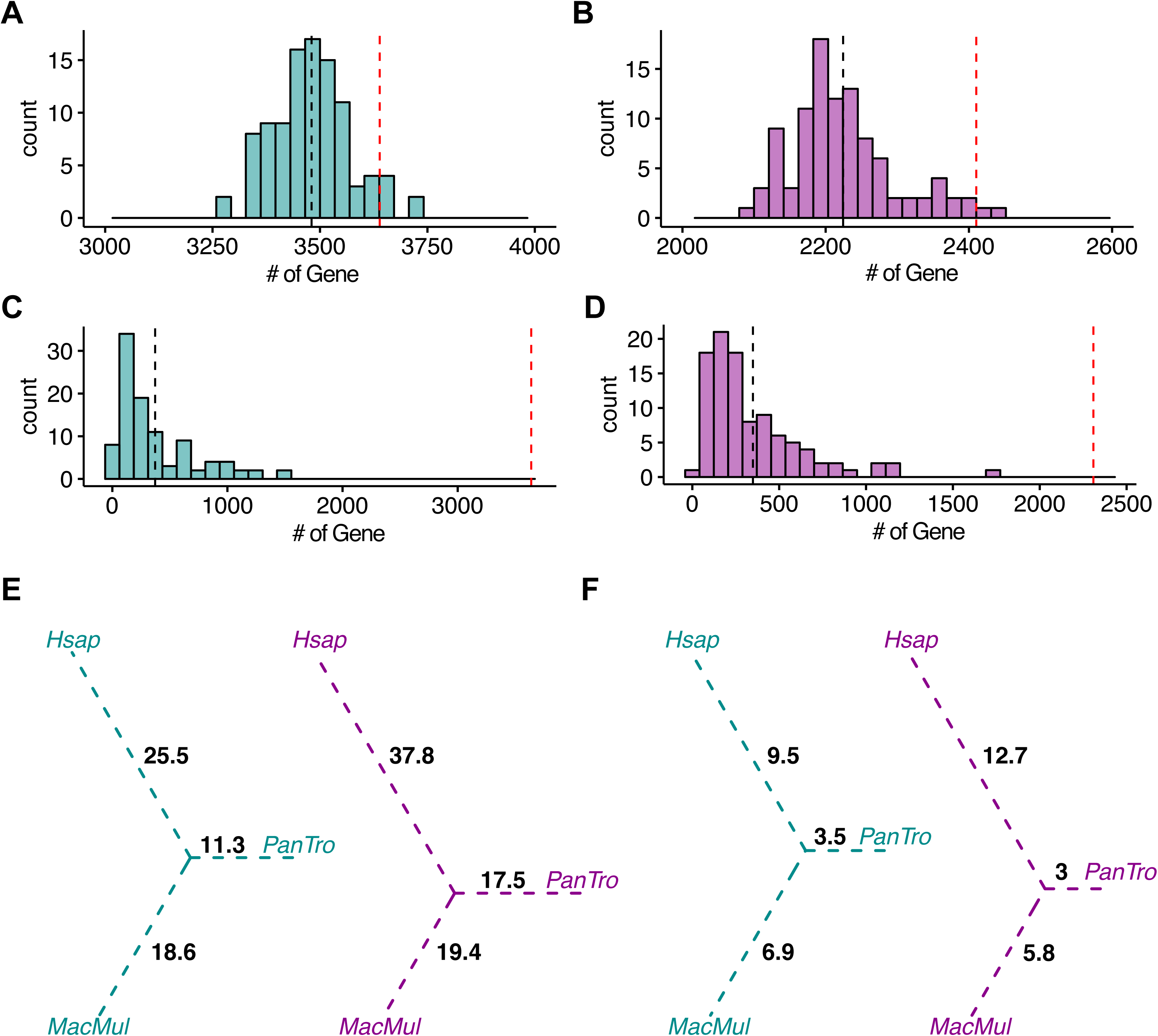
Cross-validation for differential expression analysis statistics. A-B) Leave-one-out (*LOO*) cross validation based on 100 bootstrap for (A) NeuN and (B) OLIG2. Observed number of DEGs (red dashed line) were falling in the distribution of the *LOO* DEGs based on ANOVA. C-D) Permutation analysis based on 100 permutation comparisons based on subject randomization for (C) NeuN and (D) OLIG2. Observed number of DEGs (red dashed line) were significantly different from the randomized DEGs based on ANOVA. E) Downsampled DEGs per million years based on 100 permutations. Values are calculated based on the average of species-specific DEGs. F) Downsampled DEGs per million years based on 100 permutations. Values are calculated based on the species-specific observed DEGs supported by the downsampling p-value in >90% of downsampled sets (Hsap = *Homo sapiens*, PanTro = *Pan troglodytes*, MacMul = *Macaca mulatta*).

**Figure S3.**
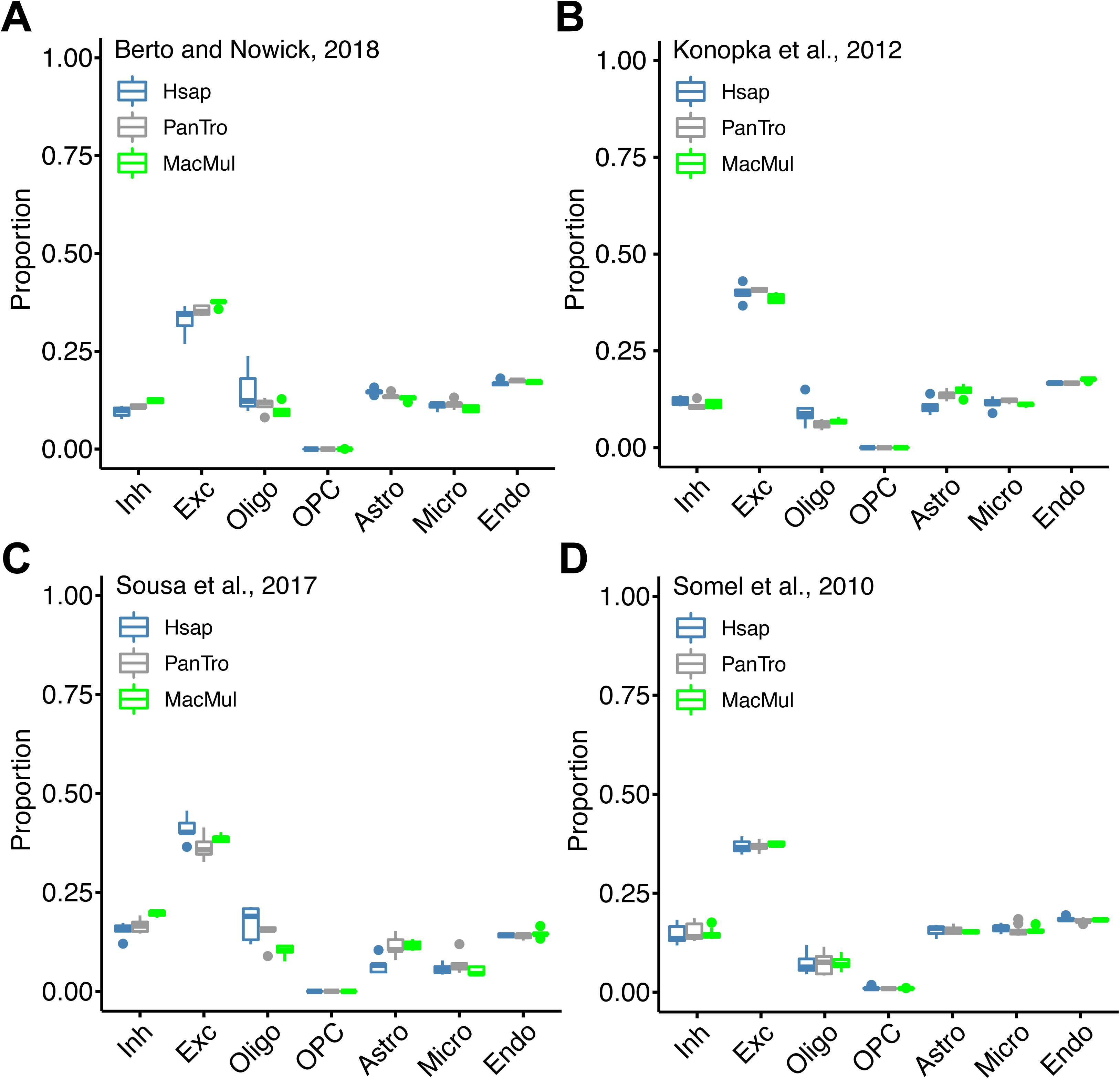
Deconvolution based on the Allen Brain Institute single nuclei data from human middle temporal gyrus (MTG) (Boldog et al., 2018). A) Primate transcriptomic meta-analysis data from Berto et al. (Berto and Nowick, 2018). B) Primate transcriptomic data from Konopka et al. (Konopka et al., 2012). C) Primate transcriptomic data from Sousa et al. (Sousa et al., 2017b). D) Primate transcriptomic data from Somel et al. (Somel et al., 2010). All data show a high proportion of excitatory neurons and sparse proportion of other cell-types (< 0.25). Y-axis represents weighted proportion. X-axis represents the cell-type identified in the single nuclei data from MTG (Boldog et al., 2018). (Hsap = *Homo sapiens*, PanTro = *Pan troglodytes*, MacMul = *Macaca mulatta*, Inh = inhibitory neurons, Exc = excitatory neurons, Oligo = oligodendrocytes, OPC = oligodendrocyte precursor cells, Astro = astrocytes, Micro = microglia, Endo = endothelial cells).

**Figure S4.**
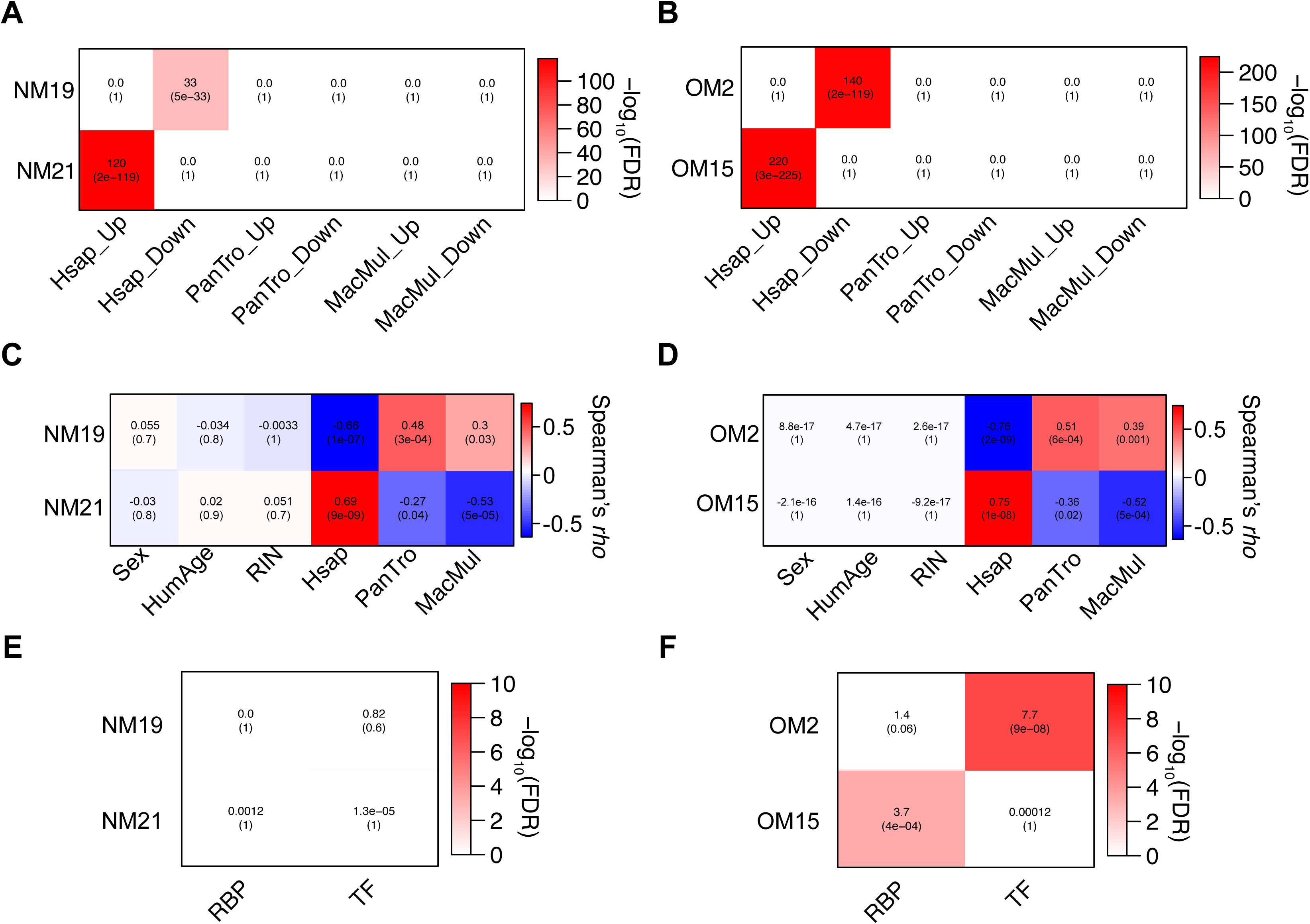
Correlations among module eigengene and associated factors. A-B) Fisher’s exact test Odds Ratios with associated FDR adjusted p-values (within parentheses) representing enrichment of species specific DEGs in (A) NeuN modules and (B) OLIG2 modules. C-D) Spearman’s rank correlations with associated p-values (within parentheses) between covariates and module eigengene of the module detected in (C) NeuN modules and (D) OLIG2 modules. (HumAge = Humanized Age, Hsap = *Homo sapiens*, PanTro = *Pan troglodytes*, MacMul = *Macaca mulatta*). None of the modules selected showed significant association with technical or biological covariates. E-F) Fisher’s exact test Odds Ratios with associated FDR adjusted p-values (within parentheses) representing enrichment of RNA-binding proteins (RBP) and transcription factors (TF) in (E) NeuN modules and (F) OLIG2 modules.

**Table S1. Demographic data.** Demographic data with life traits, RNA-seq QC metrices, technical and biological covariates.

**Table S2. Differential gene expression summary statistics for NeuN and OLIG2 data for Human, Chimpanzee, and Rhesus macaque.** All statistics for genes differentially expressed in each species, functional enrichment, and database for association with previous studies.

**Table S3. WGCNA statistics for NeuN and OLIG2 data.** All statistics for module detection, overlap with differentially expressed genes, and functional enrichment.

**Table S4. MAGMA summary statistics for NeuN and OLIG2 modules.** All statistics for GWAS enrichment in NeuN and OLIG2 modules.

**Table S5. Differential gene expression summary statistics for NeuN and OLIG2 data for Schizophrenia vs Control.** Genes differentially expressed in schizophrenia compared with controls in NeuN and OLIG2 with relative statistics.

## AKNOWLEDGMENTS

We thank the donors and their families for the tissue samples used in these studies. We thank Angela Mobley of the Flow Cytometry Facility and Vanessa Schmid of the Next Generation Sequencing Core of UT Southwestern Medical Center for technical support. G. K. is a Jon Heighten Scholar in Autism Research at UT Southwestern. This work was supported by the Uehara Memorial Foundation to N.U.; the JSPS Program for Advancing Strategic International Networks to Accelerate the Circulation of Talented Researchers (S2603) to N.U., K.T., S.B. and G.K.; the National Chimpanzee Brain Resource, NIH R24NS092988, the NIH National Center for Research Resources P51RR165 (superceded by the Office of Research Infrastructure Programs/OD P51OD11132) to T.M.P; the National Science Foundation (SBE-131719) to S.V.Y.; the James S. McDonnell Foundation 21^st^ Century Science Initiative in Understanding Human Cognition – Scholar Award to G.K.; and the NIMH (MH103517), to T.M.P., G. K., and S.V.Y. Human tissue samples were obtained from the NIH NeuroBioBank (The Harvard Brain Tissue Resource Center, funded through HHSN-271-2013-00030C; the Human Brain and Spinal Fluid Resource Center, VA West Los Angeles Healthcare Center; and the University of Miami Brain Endowment Bank) and the UT Neuropsychiatry Research Program (Dallas Brain Collection). Nonhuman primate tissue samples were obtained from Yerkes National Primate Research Center (Grant No. P51OD11132).

## Author Contributions

S.B., I.M., N.U., K.T., T.M.P., S.V.Y., and G.K. generated and analyzed the data and wrote the paper. P.C. and C.D. provided technical support. C.T. provided post-mortem human brain tissue. T.M.P., S.V.Y., and G.K. designed and supervised the study, and provided intellectual guidance. All authors discussed the results and commented on the manuscript.

## METHODS

### CONTACT FOR REAGENT AND RESOURCE SHARING

Further information and requests for resources and reagents should be directed to and will be fulfilled by the Lead Contact, Genevieve Konopka

### EXPERIMENTAL MODEL AND SUBJECT DETAILS

#### Postmortem brain samples

Human post-mortem brain samples from Brodmann area 46 were obtained from the NIH NeuroBioBank (the Harvard Brain Tissue Resource Center, the Human Brain and Spinal Fluid Resource Center, VA West Los Angeles Healthcare Center, and the University of Miami Brain Endowment Bank) and the UT Neuropsychiatry Research Program (Dallas Brain Collection) (Table S1). Nonhuman primate tissue samples were obtained from Yerkes National Primate Research Center (Table S1).

#### Nuclei extraction, FANS, and RNA isolation

Nuclei Isolation was performed as described previously (Jiang et al., 2008; Matevossian and Akbarian, 2008) with some modifications. Approximately 700 mg of frozen postmortem tissue was homogenized with lysis buffer (0.32 M sucrose, 5 mM CaCl _2_, 3 mM Mg(Ac) _2_, 0.1 mM EDTA, 10 mM Tris-HCl pH8.0, 0.1 mM PMSF, 0.1% (w/o) Triton X-100, 0.1% (w/o) NP-40, protease inhibitors (1:100) (#P8340, Sigma, St. Louis, MO), RNase inhibitors (1:200) (#AM2696, Thermo Fisher, Waltham, MA)) using a dounce homogenizer. Brain lysate was placed on a sucrose solution (1.8 M sucrose, 3mM Mg(Ac) _2_, 10mM Tris-HCl pH8.0) to create a concentration gradient. After ultracentrifuge at 24,400 rpm for 2.5 hours at 4°C, the upper layer of the supernatant was collected as the cytoplasmic fraction and set aside for RIN calculations. The pellet, which included nuclei, was resuspended with ice-cold PBS containing RNase inhibitors, and incubated with mouse Alexa488 conjugated anti-NeuN (1:200) (#MAB377X, Millipore, Billerica, MA) and rabbit Alexa555 conjugated anti-OLIG2 (1:75) (#AB9610-AF555, Millipore) antibodies with 0.5% BSA for 45 min at 4°C. Immuno-labeled nuclei were collected as NeuN-positive or OLIG2-positive populations by fluorescence-activated nuclei sorting (FANS). After sorting, gDNA and total RNA were purified from each nuclei population using a ZR-Duet DNA/RNA MiniPrep (Plus) kit (#D7003, Zymo Research, Irvine, CA) according to the manufacturer’s instruction. Total RNA was treated with DNase I after separation from gDNA. 200 ng total RNA from each sample was treated for ribosomal RNA removal using the Low Input RiboMinus Eukaryote System v2 (#A15027, ThermoFisher) according to the manufacturer’s instruction. After these purification steps, gDNA and total RNA were quantified by Qubit dsDNA HS (#Q32851, ThermoFisher) and RNA HS assay (#Q32852, ThermoFisher) kits, respectively.

#### RNA-sequencing (RNA-seq)

RNA-seq was performed as described previously (Takahashi et al., 2015) with some modifications. In order to determine the quality of the RNA from the nuclear samples, the RNA from the matched cytoplasmic fractions was extracted with the miRNeasy Mini kit (#217004, Qiagen, Hilden, Germany) according to the manufacturer’s instruction. The RNA integrity number (RIN) of total cytoplasmic RNA was quantified by an Agilent 2100 Bioanalyzer using an Agilent RNA 6000 Nano Kit (#5067-1511, Agilent, Santa Clara, CA). Samples with a total cytoplasmic RNA average RIN value of 7.5±0.16 were used for RNA-seq library preparation of the nuclear samples. For RNA-seq libraries, 50 ng of total RNA after rRNA removal was subjected to fragmentation, first and second strand syntheses, and clean up by EpiNext beads (#P1063, EpiGentek, Farmingdale, NY). Second strand cDNA was adenylated, ligated and cleaned up twice by EpiNext beads. cDNA libraries were amplified by PCR, and cleaned up twice by EpiNext beads. cDNA library quality was quantified by a 2100 Bioanalyzer using an Agilent High Sensitivity DNA Kit (#5067-4626, Agilent, Santa Clara, CA). Barcoded libraries were pooled and underwent 75 bp single-end sequencing on an Illumina NextSeq 500.

## COMPUTATIONAL METHODS

### RNA-seq mapping, QC and expression quantification

Reads from the three different primates were aligned to either the human hg19, chimpanzee PanTro4, or Rhesus macaque RheMac8 reference genome using STAR 2.5.2b (Dobin et al., 2013) with the following parameters: “*--outFilterMultimapNmax 10 --alignSJoverhangMin 10 --alignSJDBoverhangMin 1 --outFilterMismatchNmax 3 --twopassMode Basic*”. For each sample, a BAM file including mapped and unmapped reads that spanned splice junctions was produced. Secondary alignment and multi-mapped reads were further removed using in-house scripts. Only uniquely mapped reads were retained for further analyses. Quality control metrics were performed using RseqQC using the hg19 gene model provided. These steps include: number of reads after multiple-step filtering, ribosomal RNA reads depletion, and defining reads mapped to exons, UTRs, and intronic regions. Picard tool was implemented to refine the QC metrics (http://broadinstitute.github.io/picard/). CrossMap and liftOver were used to translate the non-human primate unique read coordinates into human coordinates based on hg19 (Casper et al., 2018; Zhao et al., 2014). Ensemble annotation for hg19 (version GRCh37.87) was used as reference alignment annotation and downstream quantification. Gene level expression was calculated using HTseq version 0.9.1 using intersection-strict mode by Exons (Anders et al., 2015). Counts were calculated based on protein-coding genes from the Ensemble GRCh37.87 annotation file. Orthologous genes were downloaded from Ensemble Biomart portal (Smedley et al., 2009). Orthologous genes were categorized using a high confidence score provided by ensemble and presence in known chromosomes in all three species analyzed. We removed sex chromosomes. A total of 14212 genes were considered for downstream analysis.

### Covariate adjustment and differential expression

Counts were normalized using counts per million reads (CPM) with *edgeR* package in R (Robinson et al., 2010). Normalized data were log2 scaled with an offset of 1. Genes with no reads in human, chimpanzee, or rhesus macaque samples were removed. Normalized data were assessed for effects from known biological covariates (*Gender, Age*), technical variables related to sample processing (*RIN*), and technical variables related to surrogate variation (*SVs*). Other biological and technical covariates (e.g. *Hemisphere, Pmi*) were not considered for the analysis because these were confounded with species. Non-human primates’ ages were converted to human age referring to species life traits as maximal age reached, male sexual maturity, female sexual maturity, gestation, weaning, first reproduction, number of litters, teething deciduous first and last, teething permanent first and last as proposed in (Somel et al., 2010). Traits are stored in table S1. A linear model was applied between species life traits. Human age was converted into non-human primates in R as:

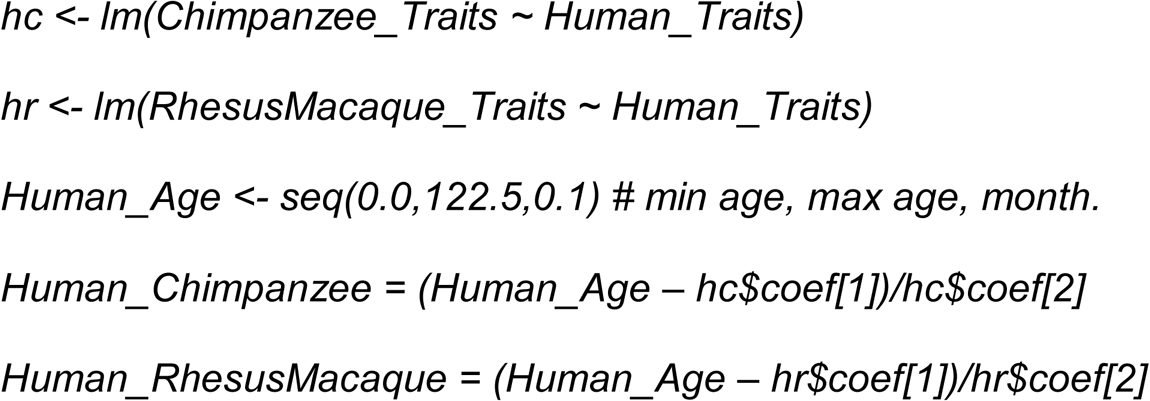

This method provided us with an accurate estimation of human age translated into non-human standard (e.g. 25 years old human corresponded to 13.2 years old chimpanzee and 8.0 years old rhesus macaque). Age was converted to categorical variables. Three groups were defined: less than 40 years old, between 40 and 60, and more than 60 years old. *SVs* were calculated using SVA in R based on a “two-step” method with 100 iterations (Leek et al., 2012). The data were adjusted for technical covariates using a linear model:

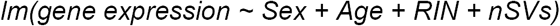

Adjusted CPM values were used for co-expression analysis and visualization. Differential expression analysis was performed in R using linear modeling. To fit our parsimony approach, we performed pairwise analysis between the three species analyzed (e.g. human – chimpanzee, human – rhesus macaque, and chimpanzee – rhesus macaque). Additionally, we performed an ANOVA based on the three species (e.g. human – chimpanzee – rhesus macaque) as following:

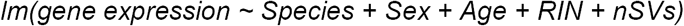

Fitting this model, we estimated log _2_ fold changes and P-values. P-values were adjusted for multiple comparisons using a Benjamini-Hochberg correction (FDR). This method was used to detect human-specific changes, chimpanzee-specific changes and rhesus macaque-specific changes using a standard cutoff of |log _2_(Fold-Change)| > 0.3 and FDR < 0.05. For example, in human, we considered specific upregulation where human showed log _2_(FC) > 0.3 and FDR < 0.05 in comparison with chimpanzee and macaque and where chimpanzee and macaque were not differentially expressed for FDR > 0.1. In addition, we considered in this paradigm the Bonferroni adjusted P-value from ANOVA of < 0.05. In contrast, for downregulated genes we consider log _2_(FC) < -0.3 and FDR < 0.05 in comparison with chimpanzee and macaque and where chimpanzee and macaque were not differentially expressed for FDR > 0.1. For the upregulated genes, we considered additional Bonferroni adjusted P-values from ANOVA of < 0.05.

### Cross-validation Analysis

To validate the robustness of our differential expression analysis, we applied a leave-one-out cross-validation by subsampling our data with *N* = # of Subject per Species –1 with # of Permutation = 100. - log _10_(Observed ANOVA) strongly correlated with -log _10_(*LOO* ANOVA), underscoring that individual subjects are not driving differential expression detection. We additionally applied a permutation method by randomizing the subjects per species 200 times and re-calculating the species-specific DEGs across subjects. The number of observed DEGs were significant different for the randomized one for both cell-type. A downsampling analysis was applied to confirm the more pronounced acceleration in OLIG2 compared with NeuN given the greater sample size for humans. Using the chimpanzee as the minimal number of subjects (NeuN = 11, OLIG2 = 10), we re-calculated the species-specific DEGs with the number of Permutation = 100. Due to the reduced sample size and high heterogeneity between and within species, the total number of species-specific DEGs was reduced. Nevertheless, this approach recapitulated the more pronounced acceleration in OLIG2, confirming the observed results based on the total number of samples.

### Co-expression network analysis

To identify modules of co-expressed genes in the RNA-seq data, we carried out weighted gene co-expression network analysis (WGCNA) (Langfelder and Horvath, 2008). Signed networks were used for both NeuN and OLIG2 data. A soft-threshold power was automatically calculated for both NeuN and OLIG2 to achieve approximate scale-free topology (R^2^>0.85). Networks were constructed with *blockwiseModules* function with biweight midcorrelation (bicor). For NeuN data, we used *corType = bicor, maxBlockSize = 10000, mergingThresh = 0.10, minCoreKME = 0.5, minKMEtoStay = 0.4, reassignThreshold = 1e-10, deepSplit = 4, detectCutHeight = 0.999, minModuleSize = 25, networkType=signed*. For OLIG2 data, we used *corType = bicor, maxBlockSize = 10000, mergingThresh = 0.10, minCoreKME = 0.5, minKMEtoStay = 0.4, reassignThreshold = 1e-10, deepSplit = 4, detectCutHeight = 0.999, minModuleSize = 35, networkType=signed*.

The modules were then determined using the dynamic tree-cutting algorithm. To ensure robustness of the observed network, we used a permutation approach recalculating the networks 200 times with permuted gene expression. Observed connectivity per gene were compared with the randomized one. None of the randomized networks showed similar connectivity, providing robustness to the network inference. We refer to this approach as permWGCNA. Additional analysis using a bootstrapping approach was performed. Briefly, we recalculated networks resampling the initial set of samples 200 times and compared the observed connectivity per gene with the randomized one. As for the permutation, none of the randomized networks showed similar connectivity. This additional test was applied to further provide robustness of the network inference.

Module sizes (25/35 respectively) were chosen to detect small modules driven by potential noise on the adjusted data. Deep split of 4 was used to split more aggressively the data and create more specific modules. Spearman’s rank correlation was used to compute module eigengene – covariates associations. A parsimony approach was used to select the modules: human-specific modules were significantly correlated with the three species but oppositely correlated between human and the non-human primates. Given the adjusted expression, covariates did not have effect on the variance explained by the gene of the detected modules. Modules were visualized based on the rank of the weight (weighted topological overlap value, WTO). Top 200 connections were selected for the visualizations. Node size was adjusted based on the *degree* (e.g. number of links). Visualization was rendered using Cytoscape (Shannon et al., 2003).

### Functional Enrichment

The functional annotation of differentially expressed and co-expressed genes was performed using ToppGene (Chen et al., 2009). Analysis was replicated using GOstats in R (Falcon and Gentleman, 2007). We used GO and KEGG databases. Pathways containing between 5 and 2000 genes were retained. Orthologous genes (14212) were used as custom background. A Benjamini-Hochberg FDR (*P* < 0.05) was applied as a multiple comparisons adjustment.

### GWAS data and enrichment

We manually compiled a set of GWAS studies for several neuropsychiatric disorders and non-brain disorders (Autism Spectrum Disorders Working Group of The Psychiatric Genomics, 2017; Bipolar et al., 2018; Davies et al., 2018; Estrada et al., 2012; Grove et al., 2017; Jansen et al., 2019; Morris et al., 2012; Okbay et al., 2016; Psychiatric, 2011; Savage et al., 2018; Schunkert et al., 2011; Wray et al., 2018). Summary statistics from the genetic data were downloaded from Psychiatric Genomics Consortium (http://www.med.unc.edu/pgc/results-and-downloads) and GIANT (https://portals.broadinstitute.org/collaboration/giant/index.php/GIANT_consortium_data_files). Gene level analysis was performed using MAGMA v1.04, which considers linkage disequilibrium between SNPs (de Leeuw et al., 2015). 1000 Genomes (EU) dataset was used as reference for linkage disequilibrium. SNPs annotation was based on the hg19 genome annotation (gencode.v19.annotation.gtf). MAGMA statistics and –log _10_(FDR) are reported in table S4 for each of the GWAS data analyzed. Brain GWAS: *ADHD = attention deficit hyperactivity disorder, ASD = autism spectrum disorders from IPSYCH (Integrative Psychiatric Research), BIP = bipolar disorder, ALZ = Alzheimer’s disease, MDD = major depressive disorder, SZ = schizophrenia*. Cognitive traits: *EduAtt = educational attainment, Intelligence = Intelligence, CognFunc = cognitive functions.* Non-Brain GWAS: *BMI = body mass index, CAD = coronary artery disease, DIAB = diabetes, HGT = height, OSTEO = osteoporosis.*

### Primate data and enrichment

Data were downloaded from respective NCBI GEO sources. Berto et al. 2018 and Konopka et al. 2012 provide species-specific differentially expressed genes within supplementary information (Berto and Nowick, 2018; Konopka et al., 2012). For the Somel et al. microarray dataset (Somel et al., 2010), raw data were downloaded and analyzed with Affy in R (Gautier et al., 2004). Degradation and quality checks were performed to the data, detecting no significant differences between the three species analyzed. We next performed a computational mask procedure using the maskBAD in R (Dannemann et al., 2012). This method developed for microarray data removed probes with binding affinity differences between species. We considered only the probesets significantly detected in at least one individual (P < 0.05). A linear model used for our data was applied to the data detecting species-specific differentially expressed genes. For Sousa et al. RNA-seq data (Sousa et al., 2017b), RPKM data were provided by the first and corresponding authors of the study. To render the data comparable with the BA46 data in our study, we used human, chimpanzee and rhesus macaque samples from dorsolateral prefrontal cortex (DFC). RPKM data were log2 scaled. Genes with RPKM = 0 in human, chimpanzee, or rhesus macaque samples were removed. A linear model was applied as used for our data to detect species-specific differentially expressed genes. Up-/Downregulated genes from these data were used for enrichment with our NeuN-/OLIG2-DEGs gene set.

### Transcription factors and RNA-binding proteins enrichment

Transcription factors list was downloaded from http://humantfs.ccbr.utoronto.ca/download.php (Lambert et al., 2018). RNA-binding proteins list was downloaded from http://rbpdb.ccbr.utoronto.ca/ (Cook et al., 2011).

### Schizophrenia cell-specific data

Differential expression analysis of cell-type expression between nuclei obtained from brain tissue derived from schizophrenia cases (SZ) and controls (CTL) was generated from GSE107638 (unpublished data). Briefly, counts were normalized using CPM with the edgeR package in R (Robinson et al., 2010). Genes with no reads in either SZ or CTL samples were further removed. Normalized data were assessed for effects from known biological covariates (Diagnosis, Age, Gender, Hemisphere), technical variables related to sample processing (RIN, Brain Bank, PMI), and technical covariates related to surrogate variation (SV). Age and PMI were converted to categorical variables (named “AgeClass” and “PmiClass”). SVs were calculates using SVA based on “be” method with 100 iterations (Leek et al., 2012). For the differential expression analysis, we used the lmTest with “robust” parameter and ebayes functions in limma package in R (Ritchie et al., 2015) fitting all the covariates reported. Significant DEGs were categorized with |log2(FC)| > 0.3 and FDR < 0.01 for both NeuN and OLIG2 cell-type SZ vs CTL analyses (table S5). Detailed information, methods and analysis are available here: https://github.com/konopkalab/Schizophrenia_CellType.

### Gene set enrichment

Gene set enrichment applied for primate DEGs as shown in Figure 1F, psychENCODE and schizophrenia cell-type DEGs as shown in Figure 4A-C and TF/RBP as shown in Figure S4 was performed using a Fisher’s exact test in R with the following parameters: alternative = “greater”, conf.level = 0.99, simulate.p.value = TRUE, B=1000. We reported Odds Ratios (OR) and Benjamini-Hochberg adjusted P-values (FDR). Enrichment was further confirmed with a hypergeometric test in R.

### Deconvolution

The human middle temporal gyrus (MTG) single nuclei RNA-seq data were downloaded from Allen Brain institute web-portal (http://celltypes.brain-map.org/rnaseq/human) (Boldog et al., 2018). Normalized data and cluster annotation were used to define cell-markers using FindAllMarkers function in Seurat (Satija et al., 2015) with the following parameters: logfc.threshold = 0.25, test.use = “wilcox”, min.pct = 0.25, only.pos = TRUE, return.thresh = 0.01, min.cells.gene = 3, min.cells.group = 3. Cell-type deconvolution was performed using MuSiC (Wang et al., 2019) with the following parameters: iter.max = 10000, nu = 1e-10, eps = 0.01, normalize=F.

### Availability of data and material

The NCBI Gene Expression Omnibus (GEO) accession number for the human data reported in this manuscript is GSE107638 (token: wlgtcayypxsjvst). Non-human primate raw data are deposited with accession number GSE123936 (token: oxyzikskbfglvwv).

### Code availability

Codes to support the DGE analysis, WGCNA, and shiny apps for data visualizations are available at https://github.com/konopkalab/Primates_CellType

